# Thermogenetic control of Ca^2+^ levels in cells and tissues

**DOI:** 10.1101/2023.03.22.533774

**Authors:** Yulia G. Ermakova, Rainer Waadt, Mehmet S. Ozturk, Matvey Roshchin, Aleksandr A. Lanin, Artem Chebotarev, Matvey Pochechuev, Valeriy Pak, Ilya Kelmanson, Daria Smolyarova, Kaya Keutler, Alexander M. Matyushenko, Christian Tischer, Pavel M. Balaban, Evgeniy S. Nikitin, Karin Schumacher, Aleksei M. Zheltikov, Robert Prevedel, Carsten Schultz, Vsevolod V. Belousov

## Abstract

Virtually all major processes in cells and tissues are regulated by calcium ions (Ca^2+^). Understanding the influence of Ca^2+^ on cell function requires technologies that allow for non-invasive manipulation of intracellular calcium levels including the formation of calcium patterns, ideally in a way that is expandable to intact organisms. The currently existing tools for optical and optogenetic Ca^2+^ manipulation are limited with respect to response time, and tissue penetration depth. Here we present **G**enetically **E**ncoded **C**alcium **Co**ntroller (**GECCO**), a system for thermogenetic Ca^2+^ manipulation based on snake TRP channels optically controlled by infrared illumination. GECCO is functional in animal and plant cells and allows studying how cells decode different profiles of Ca^2+^ signals. GECCO enabled the shaping of insulin release from β-cells, the identification of drugs that potentiate Ca^2+^-induced insulin release, and the generation of synthetic Ca^2+^ signatures in plants.

## Introduction

Calcium is one of the most evolutionary ancient signaling modalities controlling metabolism, signal transduction, and cell fate decisions in all domains of life^1^. Most major cellular functions such as cell cycle, migration, secretion, contraction, death and many others are Ca^2+^-dependent^2,3^. Ca^2+^ is a universal trigger for most types of terminally differentiated cells^4–6^.

Intracellular Ca^2+^ is regulated by a complex network of ion channels, pumps, and transporters located in membranes that maintain low cytosolic Ca^2+^ concentrations ([Ca^2+^]_i_) against the electrochemical gradient from the extracellular space and luminal compartments such as the endoplasmic reticulum (ER), lysosomes and mitochondria^3^. This underlying protein machinery permits spatio-temporally defined cytosolic Ca^2+^ patterns that are decoded by the cell as signals to change the cellular state, or function, or fate. These patterns are particularly important because elevated levels of [Ca^2+^]_i_ lead to cell death. As such, Ca^2+^ dysregulation is often associated with pathological conditions such as neurodegenerative disorders, heart disease, inflammation and diabetes^7^. Tools to generate intracellular Ca^2+^ patterns in intact cells provide a much-needed approach to manipulate and study cell signaling in a physiologically meaningful way.

Ca^2+^ transients are usually fast and local, but can also propagate over long distances^8^. There are numerous synthetic small molecules and genetically encoded fluorescent probes that allow to monitor Ca^2+^ dynamics in the living cell using fluorescence microscopy or luminometry^9,10^. However, to be able to understand the influence of Ca^2+^ on biological processes requires minimally invasive tools to manipulate Ca^2+^ at a spatial and temporal resolution that is comparable to endogenous calcium signals. Currently existing methods such as light-activatable release of Ca^2+^ from chelators or optogenetic tools often lack the dynamic range and switching stability to generate physiologically relevant calcium transients.

The first type of tools, caged Ca^2+^ compounds, offer high spatial and temporal resolution, but the system depends on the efficient delivery and loading of the caged compound, and therefore it is mostly limited to cell culture experiments^11–13^. Moreover, typical caged Ca^2+^ probes require near-UV light for activation, which induces phototoxicity, and the technique will only provide calcium transients for a short period, due to the exhaustion of the caged Ca^2+^ reservoir.

The second class of tools includes optogenetic manipulators. Channelrhodopsins conduct monovalent cations and therefore cannot be used for direct generation of Ca^2+^ transients, although the secondary increase of Ca^2+^ following plasma membrane depolarization is possible^14,15^. Recent attempts to generate genetically encoded optogenetic Ca^2+^ elevating tools led to the engineering of several light-responsive proteins. One example is a fusion of the LOV domain with the Ca^2+^ binding protein PARC^16^. A number of studies rendered store-operated calcium entry via the ORAI/STIM system, light sensitive^17–19^. Although being very selective for Ca^2+^, ORAI channels have very low conductivity. Probably due to this feature and due to its complex mechanism of action the latest version of such a tool, OptoSTIM1, demonstrated very slow dynamics of Ca^2+^ elevation in the living cells (T*_a_*_1/2_ ∼ 45 sec), and a very long (25 min) light exposure necessary to achieve activation *in vivo*^19^.

Recently, we and others developed thermogenetics, a promising neurostimulation technique that relies on a combination of temperature-sensitive ion channels and mid-infrared (IR) illumination ^20–23^. Advantages of thermogenetics stem from high conductance and fast kinetics of TRP channels used, high tissue permeability and a lack of photo-toxicity of IR light. Here, we adapt thermogenetics for the fast manipulation of Ca^2+^ levels in a variety of cells and organisms with spatial and temporal resolution not achievable before.

## Results

### A thermogenetic system for controlling intracellular Ca^2+^ levels

To thermogenetically control intracellular calcium levels and downstream physiological responses by infrared light, we employed heat-sensitive TRP channels from snakes. These channels exhibit a high prevalence for Ca^2+^ over other cations and were previously used to stimulate neurons with single cell resolution^20^. We transfected HEK293 cells with a vector encoding the TRP channel from the Texas rat snake *Elaphe obsoleta lindheimeri* (eolTRPA1)^24^ that has a temperature threshold of 38.7±0.5 °C, which is just above the normal cell culture and mammalian body temperature.

To enable optical thermogenetic stimulation, we built a custom control setup consisting of a pulsed 1 W IR laser with 1375 ± 20 nm output. (Fig. 1a, Supplementary Fig. 1). To ensure optimal positioning, the IR beam was delivered via a multimodal fiber together with a red (650 nm) guide beam visible by the naked eye and by the light detection system of the microscope (Fig. 1a,d). Shutter (On/Off) state, frequency and amplitude control provided customizable NIR pulse trains tunable with respect to the desired power, repetition frequency, and on-/offset (Fig. 1a).

**Fig. 1:**
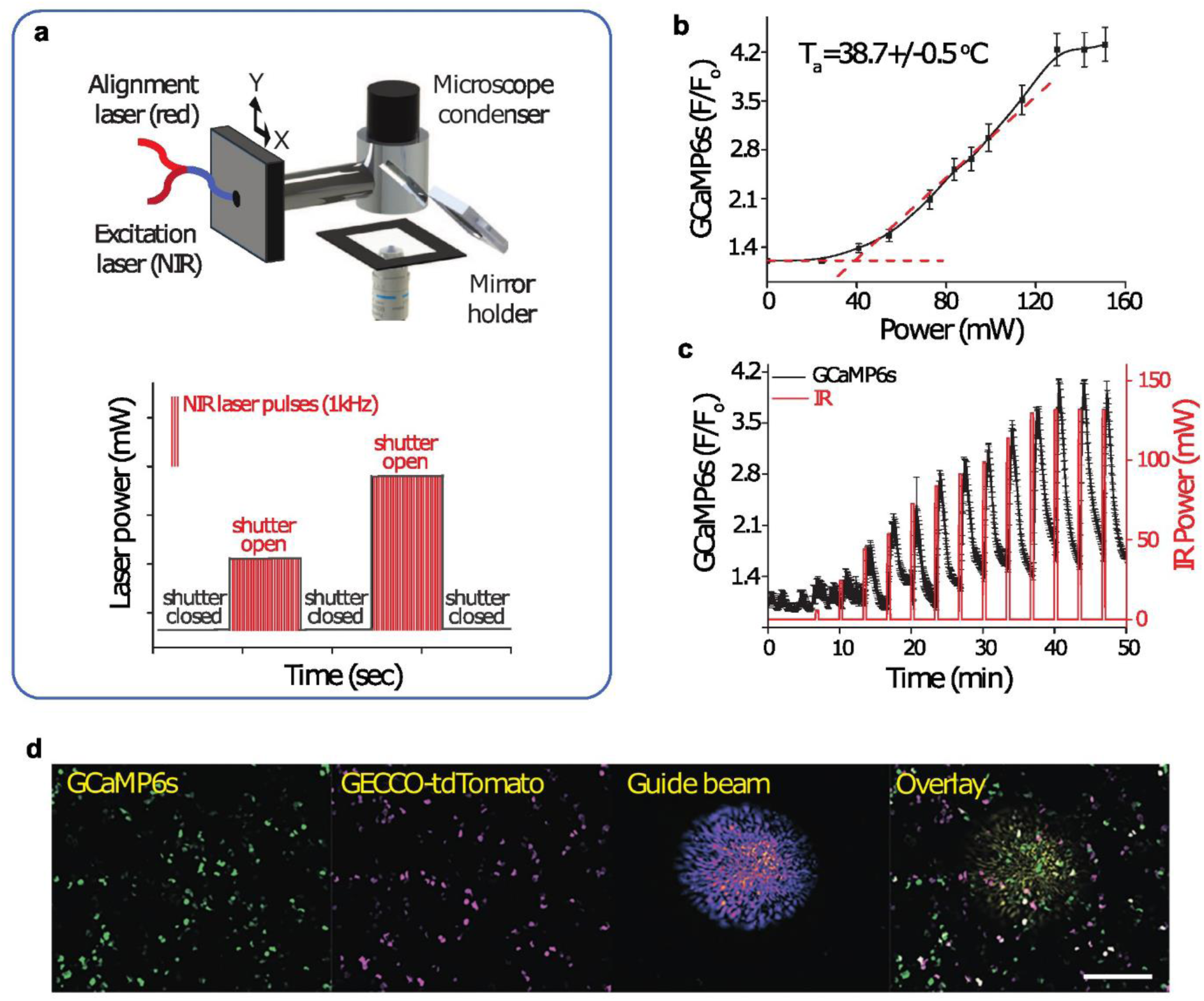
Setup for thermogenetic stimulation of cells and its parameters in mammalian cells. **a** Scheme of a mobile NIR infrared module at an inverted microscope. NIR and visible (red) lasers are optically combined to guide the NIR focal spot on the sample. The input fiber was attached using an aluminum adaptor to the condenser of an inverted fluorescent microscope. The distal end of the fiber was connected to an x-y stage to align the laser with respect to the optical axis. The lower insert illustrates a typical structure of the IR laser illumination profile used in the experiments (e.g. red profile on the panel in **C**). The profile indicates shutter opening and closing at 1 kHz pulsed NIR laser irradiation. **b** The nominal laser power-dependent eolTRPA1 temperature activation threshold (T_a_) was measured in HEK293 cells using GCaMP6s as a Ca^2+^ probe. Temperature stimulation at each point was achieved by 30 sec irradiation of 1450 nm NIR laser from the basal temperature 35°C. **c** Fluorescence intensity of GCaMP6s (black line) reflects the Ca^2+^ response in HEK293 cells to 3 to 148 mW laser stimulations (red line). **d** Fluorescent images of HEK293 cells co-expressing GGaMP6s and eolTRPA1-P2A-tdTomato (GECCO), overlaid with red laser guide beam. The scale bar is 150 μm.

To test this system, we stimulated HEK293 cells transiently co-expressing eolTRPA1-P2A-tdTomato with the calcium indicator GCaMP6s^25^ (Fig. 1b-d). Stepwise IR laser power increase from 3 to 148 mW led to progressive and proportional increases in the GCaMP6s signal amplitude (Fig. 1c). Stimulation of the cells with repeated pulses of the same intensity and duration led to the generation of highly reproducible Ca^2+^ transients (Supplementary Fig. 1). We therefore conclude that our setup allows controlling intracellular Ca^2+^ levels with high temporal resolution and good reproducibility. We termed the eolTRPA1-P2A-tdTomato system **GECCO** for **G**enetically **E**ncoded **C**alcium **Co**ntroller.

### Thermogenetic stimulation of insulin secretion

Out of various cellular processes controlled by intracellular Ca^2+^ transients, one of the most prominent is the exocytosis and secretion of hormones. An important example is the secretion of insulin from pancreatic β-cells^26^. In some forms of type II diabetes, stimulation of β-cells with glucose fails to induce adequate insulin release^27^. Therefore, new approaches to control insulin release from β-cells through tissue-penetrating light could be of great clinical importance.

It is well established that an increase of cytosolic Ca^2+^ in β-cells triggers SNARE complex formation and fusion of insulin vesicles with the plasma membrane resulting in insulin secretion^28^. Therefore, we hypothesized that by applying GECCO we may control cytosolic Ca^2+^ levels and hence insulin release (Fig. 2a).

**Fig. 2:**
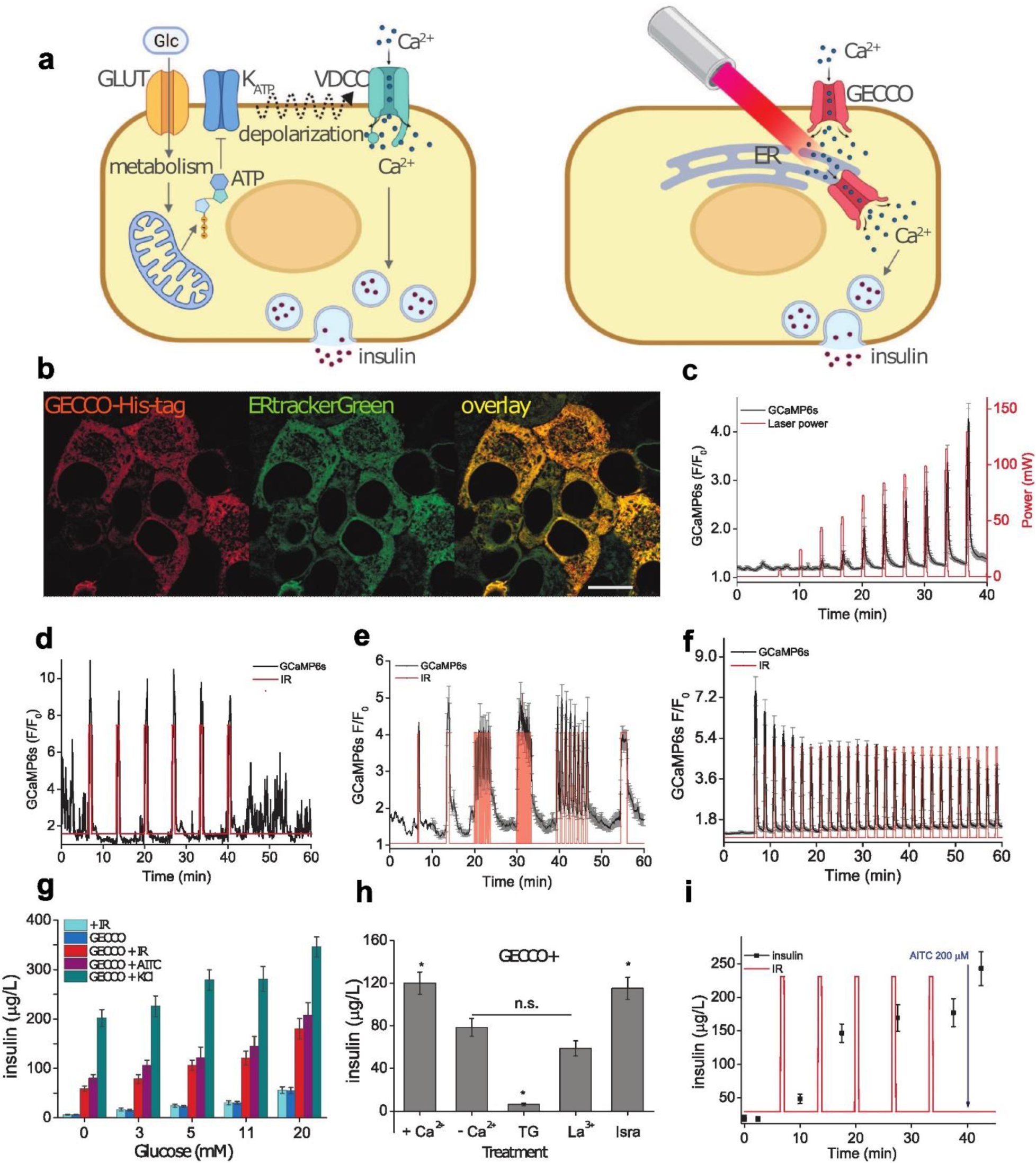
Thermogenetic control of Ca^2+^ oscillations and insulin secretion in MIN6 cells. **a** Regulatory mechanisms for insulin release in pancreatic β-cells. Left: stimulation of insulin release by extracellular glucose. Right: a proposed mechanism of thermogenetic stimulation, involving direct control of cytosolic calcium influx using GECCO located at the plasma membrane or in the ER membrane. **b** GECCO with 6xHis-tag inserted into the peripheral loop, visualized using anti-6xHis immunostaining. The cells were co-stained with ERtrackerGreen. A Pearson colocalization coefficient of 0.92 was calculated for pixels with intensities above threshold. Scale bar is 10 m. **c-f** Cytosolic Ca^2+^ profiles detected using GCaMP6s (black lines) in MIN6 cells depend on IR irradiation (red line): **c** Incremental increase in the laser power (3-148 mW). **d** Constant repeated trains (58 mW, 30 sec) for a single cell, and **f** a group of 300 cells. **e** Combination of different train frequencies and durations (3-100 sec, 0.5-10 Hz) at a constant power of 58 mW. The results are averaged from 300 cells/group from 3 replicates. **g** Insulin levels released from the MIN6 cells that were transfected with GECCO, in the presence of indicated glucose concentrations. Shown are responses to thermogenetic stimulation (58 mW, 5 trains, 30s each). Agonist stimulation was conducted using 200 μM AITC, or 10 mM KCl. **h** Insulin release from MIN6 cells under indicated conditions of calcium availability: (+Ca^2+^) medium with 2.5 mM Ca^2+^, (-Ca^2+^) nominally Ca^2+^-free medium; (TG) medium containing 5 μM thapsigargin, an inhibitor of the ER SERCA Ca^2+^ ATPase; (La^3+^) 1 mM lanthanum chloride, a plasma membrane Ca^2+^ channel blocker; (Isra) 300 nM isradipine, an inhibitor of voltage-dependent calcium channels. The stimulation profile represented in **d** was used for stimulating insulin secretion in detected using ELISA **(g and h)**. **i** Accumulation of insulin in the extracellular medium in response to GECCO stimulation. Each group of cells contained more than 6 * 10^6^ cells per three different aliquots of MIN6 cells. Laser power was 58 mW. Data represent Means ± SD.

To test our hypothesis, we used the insulinoma cell line MIN6, a widely used model for testing Ca^2+^-induced insulin release^29^. Transient expression of a 6xHis-tagged GECCO under control of the ins-2 promoter followed by immunostaining revealed a predominant localization at the ER (Fig. 2b). This location is favorable because it will permit to mimic calcium release from internal stores, a typical event in intracellular calcium signaling.

We transiently co-expressed GECCO together with GCaMP6s and performed a series of NIR-light stimulations of cells using different laser power levels and pulse train frequencies (Fig. 2c-f). In response to 30 s pulse trains of increasing laser power, GCaMP6s fluorescence in cells proportionally increased (Fig. 2c). Thermogenetic stimulation shaped the profile of the Ca^2+^ transients following the applied NIR stimulation profile, likely overriding the intrinsic depolarization induced by glucose-derived ATP levels (Fig. 2d). After the completion of GECCO stimulation trains, endogeneous Ca^2+^ oscillations were restored (Fig. 2d) indicating that the cells retained their native state.

MIN6 cells responded to NIR pulse train frequencies with reproducible amplitudes of the Ca^2+^ transients (Fig. 2e) suggesting that both kinetic properties of GECCO and the inherent capacity of the ER Ca^2+^ pumps allow precise control of cytosolic Ca^2+^ dynamics.

GECCO stimulation was able to maintain long-term stimulation with a given laser profile (Fig. 2f). Stimulation of MIN6 cells with repetitive trains of pulses (58 mW, 30 s) led to a slight gradual decrease of the Ca^2+^ amplitude, which may be attributed to a depletion of ER calcium pools.

Cultured MIN6 cells are functionally coupled via gap junctions mediated by connexins^30^. Therefore, one could expect that even random expression of GECCO in transiently transfected cells would be sufficient to activate neighboring non-transfected cells. Indeed, stimulation of GECCO-positive cells led also to Ca^2+^ transients in neighboring non-GECCO-transfected cells (Supplementary Fig. 2).

The above results show that thermogenetics can tightly control intracellular calcium levels. We next needed to investigate if different calcium signals translate into insulin secretion in β-cells.

Stimulation with repetitive IR trains led to an increase of the amount of released insulin (Fig. 2G, red bars). Application of AITC, an opener of GIRK-type ion channels, caused a similar release of insulin (Fig. 2g, violet bars). Stimulation with 10 mM KCl that induces strong depolarization of MIN6 cells caused a much higher insulin release indicating that GECCO stimulation was able to control only a part of the cellular insulin pool (Fig. 2g, green bars).

Due to its subcellular localization in MIN6 cells, GECCO likely operated with the ER Ca^2+^ pool. However, it cannot be excluded that the extracellular Ca^2+^ pool also contributed to the signal via the ORAI/STIM system or via some minor fraction of GECCO possibly residing in the plasma membrane. To test the contribution of different Ca^2+^ pools to the GECCO-mediated induction of insulin release, we performed pharmacological inhibition experiments. Application of the SERCA inhibitor thapsigargin to empty the ER Ca^2+^ pool nearly abolished insulin secretion (Fig. 2h). Elimination of Ca^2+^ from the cell culture medium, or treatment of the cells with 1 mM La^3+^, a non-specific plasma membrane Ca^2+^ channels inhibitor, also had a significant effect, probably due to opening of calcium-gated calcium channels (I_CRAC_). Inhibition of voltage-gated calcium channels (VGCCs) with isradipin had no effect on GECCO-induced insulin release.

These findings suggest that NIR-mediated insulin release in GECCO-positive MIN6 cells is predominantly regulated by ER-derived Ca^2+^ signals with a smaller contribution by calcium influx from the extracellular space. The fact that VDCCs inhibition did not affect GECCO-induced insulin release indicates that the strong cytosolic Ca^2+^ release from internal stores overrides the need for endogenous PM Ca^2+^ transport.

We next investigated the dynamics of insulin secretion in response to several sequential stimuli. As shown in Fig. 2i, the first onset of GECCO activation caused a minimal insulin release (sample 2, 5 minutes after the first stimulation). Subsequent simulations led to an increase in the insulin release (samples 3 and 4). This surprising result suggests that an occasional calcium spike will not influence insulin secretion, but that two consecutive spikes are sufficient to induce an almost full secretion response.

### Thermogenetic Ca^2+^ stimulation controls signaling pathways in β-cells

Apart from a quite direct effect on the secretion machinery, calcium ions activate a number of signaling pathways that regulate cellular functions and contribute to secretion. In this respect, the calcium-induced hydrolysis of PIP2 to DAG and IP3 leads to calcium-induced calcium release from internal stores potentiating the initial calcium effect^31^. Further, cAMP levels are known to contribute to hormone secretion^32^. We therefore investigated the interplay of GECCO-driven Ca^2+^ oscillations with DAG and cAMP formation. We employed genetically encoded biosensors for visualizing the dynamics of these intracellular messengers.

First, we imaged diacylglycerol (DAG) dynamics in MIN6 cells upon GECCO activation. DAG levels are a reliable reporter of PLC activity and is well known to stimulate PKC isoforms^33^. We employed the C1ab-GFP genetically encoded indicator^34^, which translocates to the plasma membrane upon DAG accumulation (Fig 3a-c). We co-expressed GECCO, C1ab-GFP and the red fluorescent Ca^2+^ indicator R-GECO1^35^ under control of the ins-2 promoter in MIN6 cells to perform dual parameter imaging of DAG and Ca^2+^ dynamics upon thermogenetic stimulation.

**Fig. 3:**
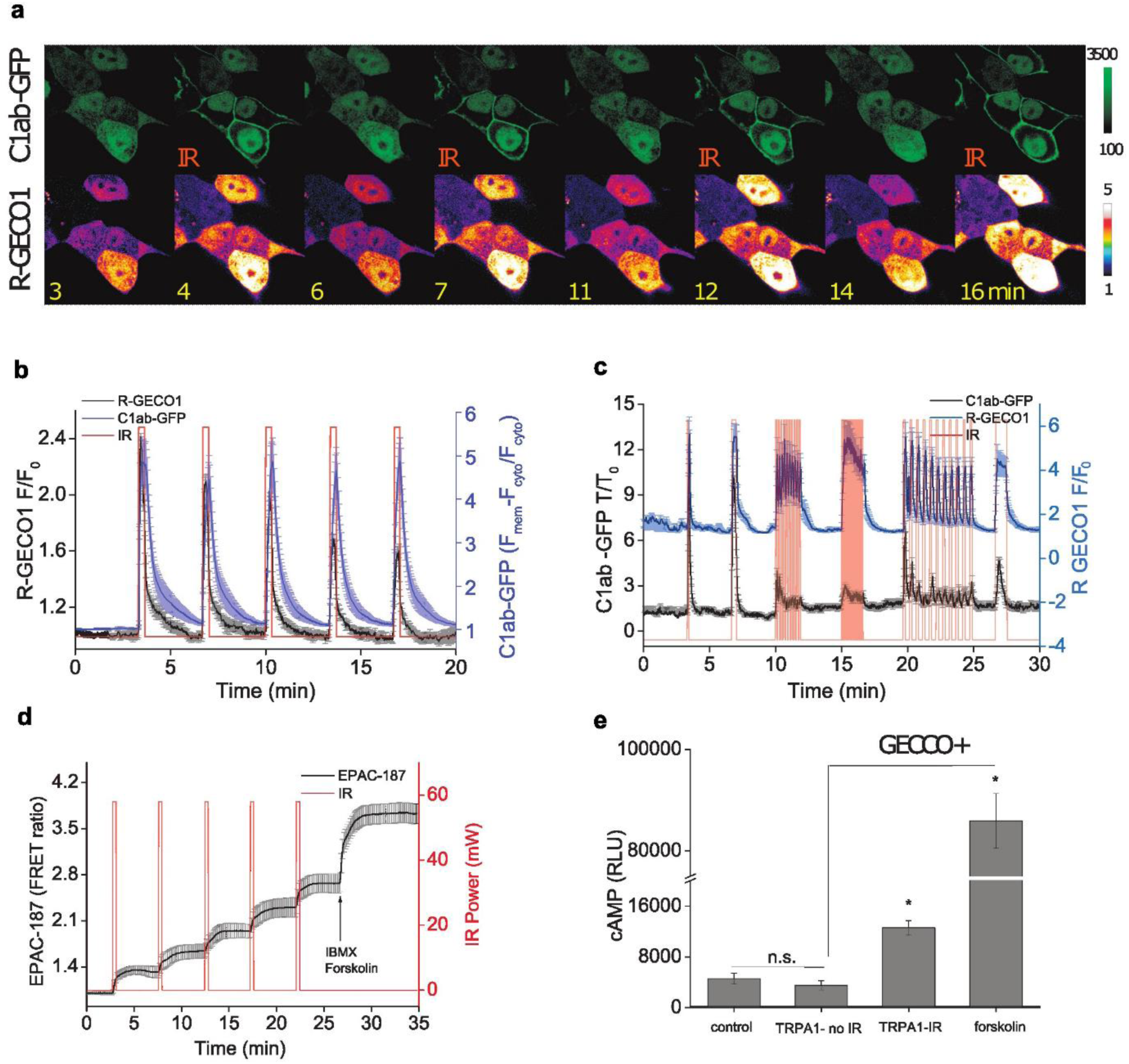
Thermogenetic stimulation alters DAG and cAMP signalling pathways in MIN6 cells. **a** Fluorescent images of cells showing simultaneous oscillations, imaged with the R-GECO1 probe, and DAG production imaged by plasma membrane translocation of C1ab-GFP. Scale bar, 10 mm. **b, c** Comparison of the normalized dynamic profiles of the cytosolic red fluorescent Ca^2+^ indicator R-GECO1 (black line), with the translocation to the plasma membrane profile the C1ab-GFP sensor to detect DAG (blue line). Detection of intracellular cAMP concentration in MIN6 cell lysates using EPAC-187 FRET-probe **d** and a bioluminescent cAMP detection system **e**.

High-speed recording demonstrated that C1ab-GFP starts translocating to the plasma membrane very rapidly, with minimal delay after R-GECO1 response, reflecting immediate PLC activation and DAG generation in response to elevated Ca^2+^ levels (Fig 3a-c). In fact, the frequency of IR stimulation was accurately reflected in C1ab-GFP transients (Fig. 3c). Albeit, the intensity of the translocation peaks decreased with increasing numbers of stimulation events. The transient nature of the DAG signal emphasizes the very rapid metabolism of DAG in MIN6 cells.

We next used the genetically encoded cAMP FRET-probe EPAC-187^36^, to detect increases in cAMP levels. Upon stimulation with repeated IR laser pulses (Fig. 3d), we observed a stepwise increase of cAMP levels. This indicates either a slow cAMP metabolism or a low off-rate of the EPAC-bases sensor. The results were confirmed with the bioluminescent cAMP-Glo kit using forskolin as a positive control (Fig. 3e).

### Harnessing thermogenetics for high-throughput screening of insulin drug candidates

Compounds that are capable of regulating insulin release are of great clinical interest. We rationalized that if we would be able to conduct GECCO stimulation in high-throughput format, it should be possible to screen for compounds that target the terminal part of the insulin release pathway between the Ca^2+^ influx and the exocytosis event.

In contrast to optogenetic tools that require visible light for activation, heat for thermogenetic activation can be delivered from non-optical generators. The most abundant devices in the modern biological laboratory, capable of providing fast heating and cooling cycles, are PCR machines, suitable as affordable tools for high-throughput thermogenetic stimulation.

In order to screen for the drugs that affect insulin release, we developed a screening system based on MIN6 cells cultured in PCR plates. Cells were subjected to heating cycles and then the insulin release was measured.

To find the optimal stimulation pattern, various PCR machine programs were tested (Fig. 4). The amount of GECCO-induced insulin released essentially depended on the stimulation pattern (Fig. 4b). Remarkably, we observed an increase of insulin secretion upon elevation of the stimulation frequency up to 5 Hz. This may indirectly evidence the presence of a mechanism enabling β-cells to integrate multiple Ca^2+^ waves determining the physiologically relevant output (level of insulin released).

**Fig. 4:**
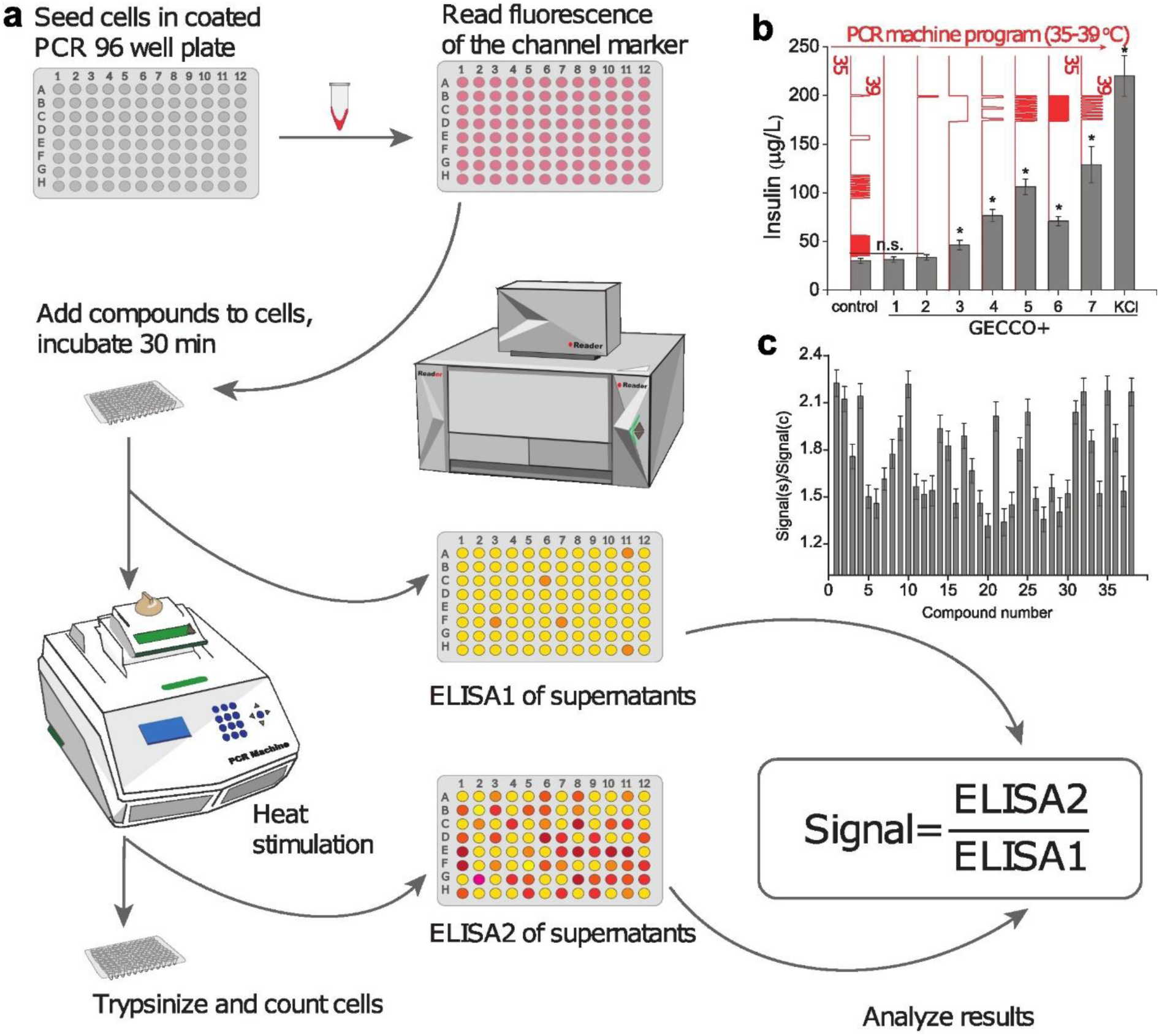
Drug screening using high-throughput thermogenetic stimulation in a PCR machine. **a** Experimental design. Cells were seeded in poly-D-lysine coated 96-well PCR plates and transfected with GECCO-P2A-tdTomato. The transfection level was measured using a fluorescent plate reader (Tecan Infinite Pro). Each well was supplemented with a selected drug (1 drug/1 well in quadruplicates) and incubated for 30 min at 35 °C following by sampling of cell media (10 ml) for ELISA1 and replacement of 10 ml fresh media. Then the plates were placed in a PCR machine and stimulated by heating-cooling cycles (39 °C – 30 s, 35 °C – 3 min, 40 cycles), and 10 ml of cell medium from each well was sampled for ELISA2. Each drug was assessed in 3 replicates. **b** ELISA analysis of insulin released by MIN6-GECCO cells into the medium depending on the shape of the thermogenetic stimulation programmed by the PCR machine. **c** The results of the thermogenetic stimulation of insulin release in MIN6 cells, pretreated with FDA-approved drug panel (see Supplementary Tables 1 and 2 for details).

Next we tested a library of FDA-approved drugs including more than 800 compounds for their ability to affect GECCO-induced insulin release. Cells were pre-incubated with drugs for 30 min before the stimulation and subjected to the optimal stimulation protocol.

Drug responses were divided into two groups. First group of drugs induced insulin release regardless the thermogenetic stimulation (Supplementary Table 1). But we have identified also a group of compounds that altered only the GECCO-induced insulin release, but did not affect the insulin release in the absence of thermogenetic stimulation (Supplementary Table 2). These results confirm the utility of GECCO-based high-throughput screening systems for drugs affecting Ca^2+^-dependent processes in cells.

### Thermogenetic control of Ca^2+^ in plants

Application of optogenetic tools to plant systems has been limited due to a high optical density of plants^37,38^, and co-activation of the host photoreceptors. Moreover, ambient light required for plant growth obviously induces undesired activation of many optogenetic tools. Therefore, application of the thermogenetics-based GECCO system in plants could provide a suitable alternative ^39–42^.

Lab-grown plants are cultivated at much lower temperatures than those typical for mammalian cells. Therefore, we employed the caTRPA1 ion channel from rattle snake that exhibits a temperature threshold of 28 °C^20,24^. We named this system GreenGECCO to highlight its utility for Ca^2+^ control in plants.

After initial tests demonstrating that GreenGECCO is able to evoke Ca^2+^ transients in *Nicotiana benthamiana* leaves under NIR illumination (Supplementary Fig. 3), we transformed transgenic *Arabidopsis thaliana* plants, stably expressing the Ca^2+^ probe R-GECO1^43^, with caTRPA1-GSL-mTurquoise (GreenGECCO). GreenGECCO expression in *Arabidopsis* induced signs of toxicity when plants were grown at 22°C. To avoid such toxic effects we cultivated the plants at 16-18 °C. We stimulated the root tip of 4-day-old Arabidopsis seedlings with increasing intensities of NIR pulses, and analyzed the cytosolic Ca^2+^ signals spreading from the root tip. With increasing NIR pulse power, the amplitude of the calcium response increased (Fig. 5a). Laser power above 90 mW induced long-distance propagating Ca^2+^ waves (Fig 5a). In plants expressing the control construct R-GECO1-mTurquoise^44^ alone, such stimulations did not cause any changes in intracellular Ca^2+^, indicating that the signals obtained were specifically caused by GreenGECCO stimulation (Supplementary Fig. 4a).

**Fig. 5:**
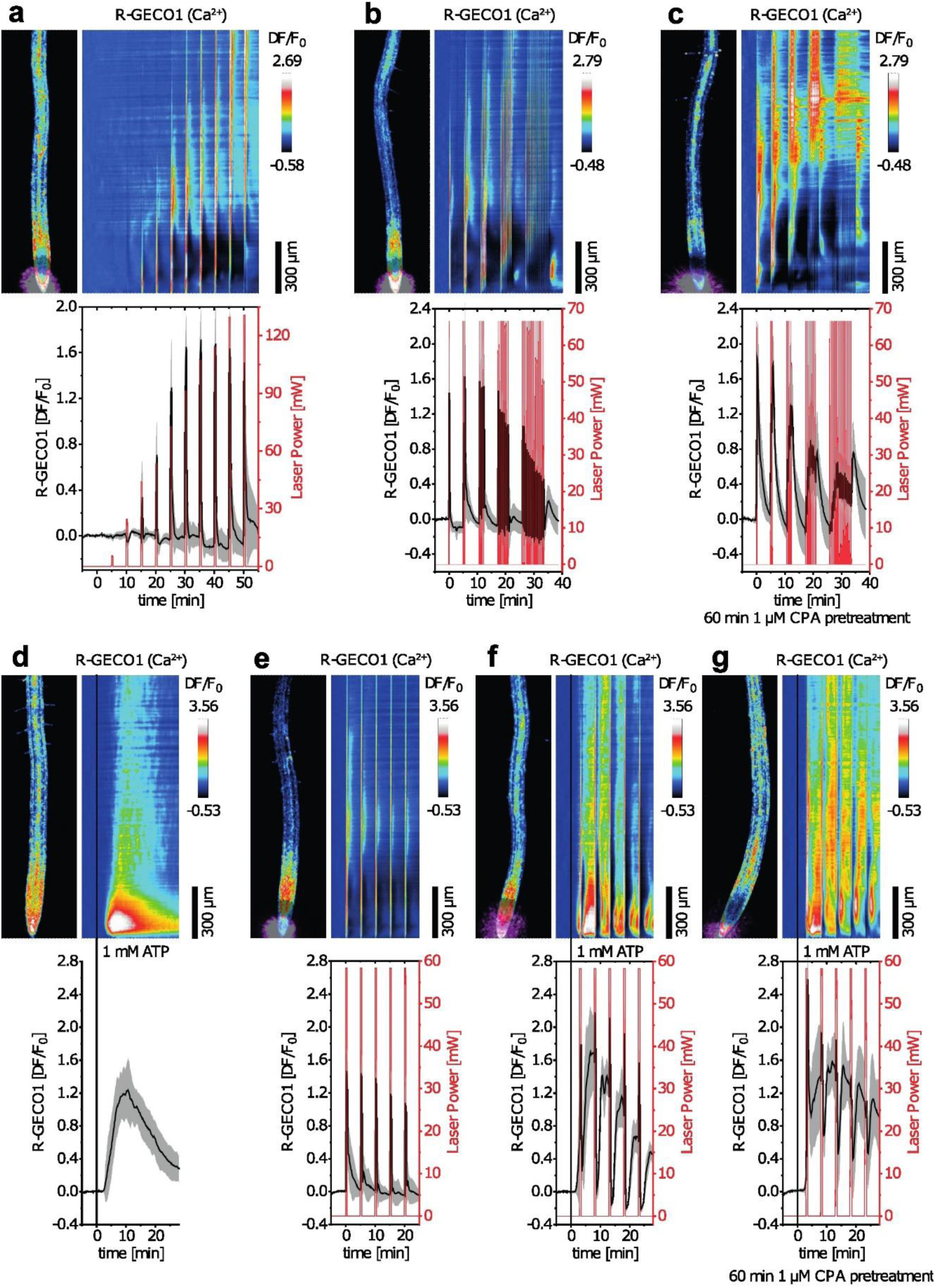
Thermogenetic control of cytosolic Ca^2+^ levels in Arabidopsis. **a-g** Four-day-old Arabidopsis seedlings, expressing caTRPA1-GSL-mTurquoise (GreenGECCO) in the R-GECO1 Ca^2+^ probe background, were thermogenetically stimulated at the root tip. The left images indicate the respective R-GECO1 fluorescence intensity at t = −5 min, with the purple spot representing the region of NIR laser stimulation. The right images indicate normalized average spatiotemporal response profiles of R-GECO1 (color scale bar on the right) along the root axis with the same time scale as in the graph below (frame rate 10 min^−1^). The graphs indicate normalized average R-GECO1 responses (black lines), calculated from whole image mean values, and the applied NIR laser power (red lines). **a**, Responses to 30 sec pulses of increasing NIR laser power with 4.5 min gaps between each pulse (means ± SD, n = 6). **b-c**, Responses to increasing numbers (1, 2, 4, 8, 16) of 6 sec 67 mW pulses with 30 sec of recovery after each pulse without **b** and with **c** 1 h pretreatment with 1 µM of the ER Ca^2+^-ATPase inhibitor cyclopiazonic acid (CPA) (means ± SD, n = 5). **d-g** Combined thermogenetic and 1 mM extracellular ATP stimulations. **d** Stimulation with 1 mM extracellular ATP applied at t = 0 min. **e** Thermogenetic stimulations using five consecutive 30 sec pulses of 58 mW NIR laser power with 5 min intervals. **f** 1 mM extracellular ATP stimulation at t = 0 min and the five-pulse thermogenetic stimulation applied at t = 3 min. **g** As in **f**, but 1 h after pretreatment with 1 µM CPA (means ± SD, n = 6).

Application of various NIR pulse patterns revealed a phase locked Ca^2+^ response in the cytosol (Fig. 5b), indicating that calcium signaling in plants can operate at various frequencies. Interestingly, long-distance Ca^2+^ signals became more apparent with increasing NIR frequencies (Fig. 5b).

Confocal microscopy suggested an ER localization of GreenGECCO-GSL-mTurquoise in leaf epidermis and root tip cells (Supplementary Fig. 4b), indicating that the leading calcium source in thermogenetic stimulation in plants is likely the ER. Inhibition of extracellular calcium influx with 90 min 1 mM LaCl_3_ pretreatments of seedlings resulted in only slightly decreased intensities of calcium spikes (Supplementary Fig. 4b).

Interestingly, inhibition of ER-type Ca^2+^-ATPases (ECAs) with 1 µM cyclopiazonic acid (CPA) resulted in prolonged Ca^2+^-transients, indicating that these Ca^2+^ pumps are actively involved in shaping of rapid Ca^2+^ signals by pumping cytosolic Ca^2+^ into the ER lumen (Fig. 5c).

Naturally occurring Ca^2+^ waves in plants are able to propagate over long distances between the plant organs^8^. GreenGECCO-induced Ca^2+^ waves excited in either the hypocotyl or the root propagated from their origin to neighboring tissues (Supplementary Notes, Supplementary Fig. 6d,e).

In plants, extracellular ATP is a strong elicitor for cytosolic Ca^2+^ elevations^45,46^. In our experiments, addition of 1 mM ATP to Arabidopsis root tips induced “gaussian-like” calcium signatures (Fig. 5d), as observed previously^47^. Here, we asked whether and how thermogenetically induced Ca^2+^ spikes interfere with such ATP-induced Ca^2+^ signals. We combined thermogenetic stimulations with ATP treatment (Fig. 5e,f). This resulted not only in the superposition of both signal profiles, but also induced sharp decays of cytosolic calcium levels after the termination of each NIR stimulation. This indicates that GECCO-evoked cytosolic Ca^2+^ increases in plants and activate rapid and efficient Ca^2+^ export systems and that constant ATP stimulation of *Arabidopsis* roots maintains a characteristic response profile. One of the rapid Ca^2+^-export systems might be associated with ECA-type Ca^2+^-ATPases, as CPA inhibition of these pumps attenuated the observed Ca^2+^ decays (Fig. 5g).

Together, these results demonstrate the applicability of GreenGECCO as a tool to generate synthetic Ca^2+^ signals in plants with high spatiotemporal resolution.

## Discussion

In summary, GECCO meets the need for a genetically encoded, robust, high-speed, high-conductance system to operate Ca^2+^ without phototoxic effects. Based on the TRPA1 channel with high conductance and great calcium preference, GECCO can be activated by infrared wavelengths or, in principle, any other source of heat radiation. The precise switching temperature ensures perfect control without the danger of over-heating the sample. Once expressed, GECCO directly conducts Ca^2+^ without a need of additional synthetic small molecules or cofactors.

TRP channels have lower ion selectivity compared to, for example, Opto-STIM. While ORAI channels are exclusively selective for Ca^2+^, TRP channels can conduct not only calcium, but also monovalent cations such as sodium and potassium. However, they conduct preferably Ca^2+^ because the Ca^2+^ concentration gradients are much steeper across cellular membranes separating the cytosol (nM for Ca^2+^) from the ER lumen or the extracellular space (mM Ca^2+^ in both compartments).

Using GECCO, we were able to control insulin secretion from β-cells. There were earlier attempts to control insulin secretion using optogenetics. However, optogenetic stimulation increased Ca^2+^ indirectly, via depolarization of the plasma membrane activating voltage-dependent Ca^2+^ influx^48,49^. As a result, it was challenging to achieve dose dependent Ca^2+^ influx and insulin release. In the present study, we demonstrate that GECCO controls insulin release from β-cells in a Ca^2+^ dependent manner. A big advantage of this stimulation scheme is its simplicity. Ca^2+^ elevation in the cytosol is the last step of the insulin release cascade directly preceding the fusion of insulin granules with the plasma membrane. Therefore, the molecular path from GECCO stimulation and insulin release is very short and easy to control.

As a Ca^2+^ store, the endoplasmic reticulum in β-cells seem to play a dual role. It can contribute to a cytosolic Ca^2+^ levels via the mechanism of Ca^2+^-induced Ca^2+^ release^50^. On the other hand, the ER serves as a sink responsible for the removal of the cytosolic Ca^2+^ via SERCA. However, to date it is unclear if Ca^2+^release from the ER is sufficient to induce insulin secretion. Our results demonstrate that, i) in principle, the capacity of the ER Ca^2+^ pool is sufficient to induce insulin secretion, as inhibition of Ca^2+^ entry via the plasma membrane by La^3+^ ions only partially reduces insulin release, and ii) at least in the conditions of GECCO stimulation, the ER Ca^2+^ pool plays the main role in the secretion, as thapsigargin completely abolished the insulin secretion.

Another important observation is that in response to a single stimulation, β-cells secrete only small amount of insulin, whereas repetitive stimuli lead to much stronger release (Fig. 2i). Similar conclusions could be made from the experiment of thermogenetic stimulation using a PCR machine to pulse the temperature. Single cycles resulted in much less insulin release to the media compared to trains of the stimuli (Supplementary Fig. 4b). Moreover, three sequential clearly separated short heating-cooling cycles produced higher levels of insulin release compared to a single continuous heating cycle of the same duration (compare patterns 3 and 4 on the Supplementary Fig. 4b). Likely, β-cells resist random spontaneous Ca^2+^ increases and require sequential raises of sufficient amplitude to induce insulin secretion.

Calcium is an essential regulator and messenger in all domains of life, including plants where Ca^2+^ coordinates many activities such as growth, fertilization and immune responses ^51^. Although optogenetic Ca^2+^ modulators are regularly employed in animal cells, such tools for specifically controlling cytosolic Ca^2+^ levels in plant cells are currently not available^40,52^. Optogenetics in plants has several disadvantages. (i) Plant tissues are poorly permeable for visible light wavelengths making optogenetic activation complicated. (ii) Visible light activates not only optogenetic systems in plants, but also endogenous plant photoreceptors. (iii) Ambient light required for plant growth induces undesired activation of the optogenetic tools. We were able to induce Ca^2+^ transients in transgenic plants expressing the GECCO system. Using thermogenetics, we were able to overcome these hurdles. Thermostimulation of GreenGECCO expressing *Arabidopsis* plants allowed very precise programming of synthetic Ca^2+^ signatures and the induction of long-distance Ca^2+^ waves (Fig 5, Supplementary Fig. 6) opening new avenues for studying the effects of distinct Ca^2+^ signatures on plant physiology and development.

Thermogenetic calcium stimulation holds a great promise not only for fundamental research, but also for translational medicine. Unlike optogenetics that is mostly used to activate neurons, GECCO can efficiently modulate the activity of various cell types by modulating intracellular Ca^2+^. The major limitation of current optogenetic tools, like ChRs, in translational medicine is their non-human origin that inevitably would induce an immune response in the human organism leading to elimination of cells expressing the channelrhodopsin or other similar molecules. Humans have their own TRP channels that open a wide avenue for potential therapeutic modalities in various fields of medicine in the future.

## Acknowledgments

We thank Christian Waadt for developing the plant GEFI analyzer Fiji plugin, the EMBL Chemical Biology Core Facility for providing a library of FDA approved drugs. R.W. was supported by the Deutsche Forschungsgemeinscahft (WA 3768/1-1). R.P. and C.S. were supported by the European Molecular Biology Laboratory (EMBL), M.S.O. acknowledges a fellowship under the Marie Skłodowska-Curie Actions COFUND (grant agreement number 664726). YGE was supported by FEBS STF, DAAD LTF, S. Shpiz stipend; V.V.B. was supported by Priority 2030 program from the Ministry of Science and Higher Education of the Russian Federation. C.S. acknowledges funding from TRR186/Mercator Fellowship by the Deutsche Forschungsgemeinschaft,

## Methods

### Plasmids and genes

All constructs for mammalian cells were cloned with NEBuilder HiFi DNA Assembly Master Mix (#E2621L, NEB) from PCR fragments obtained with Q5 High-Fidelity 2X Master Mix (#M0492L, NEB). eolTRPA1 coding sequence from Texas Rat snake (Elaphe (Pantherophis) obsoletus lindheimeri, GenBank: GU562966.1), with P2A-tdTomato tag, was cloned under ins2 promoter from pGL410_Ins2500 (#49058, Addgene) and inserted into pCAGGS from pCAGGS-PSD95-GCaMP2 (#18931, Addgene) vector for specific expression in β-cells. The correctness of all constructs was verified by sequencing (eurofinsgenomics.eu).

### Cell culture. HEK293

Human embryonic kidney cells were received from ATCC stocks of EMBL Heidelberg in C. Schultz lab, and cultured at 37°C and 8.0% CO_2_ in high-glucose DMEM (#41965-039, Life Technologies) supplemented with 10% Fetal Bovine Serum (10270098, Life Technologies), penicillin–streptomycin (Pen Strep, 100 U/mL, #15140122, Life Technologies). Cells were split by 3 min incubation at 37°C with 1x TryplE, centrifugation at 800 *g* for 5 min, and 1:10 seeding to a new T25 flask.

### MIN6

Low passage mouse insulinoma-derived cells, the MIN6 cell line (passage 27-39) was initially developed and provided as a kind gift from J.-I. Miyazaki. MIN6 cells were cultured at 37°C and 8.0% CO_2_ in high-glucose DMEM (#41965-039, Life Technologies) with 15% FBS (10270098, Life Technologies), penicillin–streptomycin (Pen-Strep, 100 U/mL, #15140122, Life Technologies) and 2-mercaptoethanol (70 μM solution in PBS, #P07-05100, PAN-Biotech) was added freshly to the cell medium just before use.

For microscopy studies, cells were seeded in 8-well Lab-Tek (#155411, Thermo Scientific Nunc) or 24-well glass-bottom (#P24-1.5H-N, Cellvis) dishes to reach 20–30% confluence. For fluorescent plate reader experiments cells were seeded in flat bottom clear, white polystyrene 96 well glass-bottom plates (#CLS3610, Corning). After 48 h MIN6 cells were co-transfected with a plasmid of interest. The transfection mix was prepared in Opti-MEM (50 μL/well, 31985-070, Life Technologies) with Lipofectamine 2000 transfection reagent (0.5 μL per well, #11668030, Life Technologies, according to the manufacturer’s protocol) with 160 ng of total plasmid DNA (ratio of the transfection reagent:DNA = 3:1) and added to each well of an 8-well Lab-Tek microscope dish with cells preloaded with 300 μL fresh Opti-MEM.

For 96-well plates we prepared 100 uL of Opti-MEM per well transfection mix containing 50 ng of the plasmid DNA and 0.17 μL of the Lipofectamine 2000 and replaced the cell medium with the transfection mix. All fluorescent probes were transfected into four individual plates in quadruplicates per plate.

After 12 h of incubation at 37°C and 8.0% CO_2_, cells were rinsed twice with Opti-MEM and the medium was exchanged for the culture medium, followed by incubation for another 24-48 h (HEK293) or 48-72 h (MIN6).

For in vitro biochemical assays cells were cultured on d60 mm /d35 mm dishes (#150288/153066, Nunc, Thermo Scientific) in standard conditions without transfection.

### Insulin ELISA

Enzyme-linked immunosorbent assay (ELISA) for the quantification of insulin content in the cell medium of MIN6 and mouse primary cells was performed in 96-well format (#10-1247-01 Mercodia AB, Uppsala). Insulin levels were normalized to the number of cells in the dish or well. The cells were counted with TC20™ Automated Cell Counter (#145-0102, Biorad). Cells were detached by TrypLE (#12604013, Thermo Scientific) treatment for 5 min at 37°C and 8.0% CO_2_. The total protein content in the MIN6 cell medium was determined using a BCA standard normalization (Pierce TM BCA Protein assay kit, Thermo Scientifiс), and total DNA quantification was conducted using a General DNA Quantification Kit (# ab156902, Abcam). Accumulation of DNA or total protein content increase without change in insulin release in the cell medium was considered as a toxicity marker.

For correlation analysis of calcium spikes during microscopy and insulin secretion in one sample, we took samples of the medium (10 µL/sample, 4% of the medium, replacing for the same aliquot of inoculated media).

Prior to the experiments, cells were carefully washed twice from serum with Krebs-Ringer solution supplied with 0-22 mM glucose. After 30 min a fresh aliquot of Krebs-Ringer solution was applied for insulin secretion experiments. Samples were collected after incubation for 40 min at 37°C. Experiments for the determination of insulin content were performed in quadruplicates per condition.

In the experiment relative to Fig. 2i, after initial 20 min incubation of the cells in a fresh medium, cells were stimulated with five sequential IR trains. After each train 5% of the culture medium was sampled for ELISA assays and replaced with fresh medium. Medium sampled for insulin quantification before the first stimulation was used to measure the baseline level. After the last NIR pulse cells were treated with 200 μM AITC. The amount of insulin released was normalized to the total cell protein concentration and DNA content.

### cAMP detection

Detection of the cAMP was performed using a commercial luciferase-based kit (#V1501, Promega) in 384-well format using cell lysates according to the manufacturer’s protocol. The bioluminescent signal was detected with a Tecan Infinite M1000 Pro reader in 384-well Armadillo Plates (BC-3384, Thermo Scientific).

### ER labeling and ICC staining

For ER labeling we used ERTrackerGreen (BODIPY FL Glibenclamide, ThermoFisher Scientific, # E34251) according to the manufacturer’s protocol (#MP 12353). Excitation/Emission of ER-Tracker Green was 504/511 nm, making it available for imaging using GCaMP6s microscope settings.

For ICC, cells in 8-well Lab-Tek plates were placed for 20 min at room temperature (RT) in PFA/SEM buffer (4% paraformaldehyde, 0.12 M sucrose, 3 mM EGTA, and 2.5 mM MgCl_2_ in PBS), washed with PBS (300 µL/well) and aldehyde groups were quenched with 50 mM NH_4_Cl for 10 min at RT. Cells were permeabilized with 0.1% Triton X-100 for 10 min at RT. Permeabilized cells were washed in PBS, 3x 5 min at RT (300 µL/well) and blocked with 5% BSA with 2% donkey serum in PBS for 30 min at RT. 6x-His Epitope Tag Monoclonal Antibody (#MA1-21315, Thermofisher Scientific; dilution 1:500) was used for His-tag staining of the TRPA1 channel in fixed cells for 1 h at RT. After 3x 5 min 300 µL/well PBS washes at RT, donkey anti-Mouse, Alexa Fluor Plus 594 Anti-Mouse Secondary Antibody (#A32744, Thermofisher Scientific) at a dilution of 1:500 was used as a secondary antibody for 40 minutes staining at RT. Final samples were washed 3x with PBS and used for fluorescent imaging at a FluoView 1200 confocal laser scanning microscope at RT in 300 µL PBS.

### Confocal microscopy of mammalian cells

Before starting the imaging, cells were preincubated for at least 20 min at 37°C in 250 μL imaging buffer containing115 mM NaCl, 1.2 mM CaCl_2_, 1.2 mM MgCl_2_, 1.2 mM K_2_HPO_4_, 18 mM HEPES, 0-22 mM glucose adjusted to pH 7.2. Confocal imaging was performed at 37 °C using a dual scanning FluoView 1200 confocal laser scanning microscope equipped with 20x UPLS APO (NA 0.75, air) and 60x Plan-APON (NA 1.4, oil) objectives and a Hamamatsu C9100-50 EM CCD camera. Single-FP based probes (GCaMP6s, C1ab-GFP, R-GECO1) were imaged in one fluorescent channel: blue for C1ab-GFP and GCaMP6s with 488 nm laser line (1.0-3.0 %), and red for R-GECO1 with 561 nm excitation laser, 580-630 nm bandpass emission filter, HV 500-700 mV, pinhole 50-150 microns, 4 sec between frames. The FRET probe EPAC was excited with a 405nm laser line with the donor emission collected from 460 to 510 nm and the acceptor emission collected from 515 to 540 nm, pinhole 300 microns, 10 sec per frame.

For all reagents and compounds, the used concentrations are shown in the corresponding figures and legends, and all stocks were prepared as 100x concentration.

### Analysis of cell imaging data

Imaging analysis was performed with the Fiji (ImageJ) FluoQ^53^ macro. Fluorescent intensities of individual cells were extracted after segmentation, background subtraction and automatically accumulated as mean ± SD groups. At least 300 individual cells in 4-6 independent experiments per condition were selected. A ratiometric image was calculated as the mean of all segmented cells followed over time, normalized to the minimum value and plotted. All data groups passed normal distribution tests. For simple pair-wise comparisons a paired two-tailed Student’s t-test was performed. For multiple comparisons analysis of variance (ANOVA), a comparison followed with Bonferroni correction and Rodger’s method post-hoc test was used. Data were considered to be statistically significant when final p < 0.05. The resulting data were statistically analyzed and plotted with Origin 8.6 Pro followed by a figure assembly in Adobe Illustrator 11.

### Infrared stimulation setup

The IR optical module was designed to stimulate cells by optical heating, and used for cell cultures and *Arabidopsis* experiments where large groups of cells should be heated. Our custom design module was tailored to fit into the condenser of a conventional microscope (FluoView 1200-1000, Olympus) but in principle, it can be adapted to fit any microscope inverted microscope (Supplementary Fig. 1).

The module consists of an IR laser (1375nm laser +/- 20nm, 4PN-117, Seminex, USA) for thermal stimulation which was driven by a current driver (4320, 20A, Arroyo Instruments) and by maintaining the stability of laser power with a temperature control unit (ARR 5305, Arroyo Instruments). A function generator was used to trigger the NIR laser and form a periodic On-Off states for stimulation at a desired power and duration.

To aid in alignment, the IR laser was combined with a visible laser (650nm, 5mW, Fiber Fault Locator laser pen) via a 2×1 fiber coupler (TM105R2F1B, Thorlabs) (Supplementary Fig. 1a). The distal end of the fiber coupler was connected to an X-Y stage for lateral adjustment. The lasers were then relayed onto the sample with a 4F imaging system by reflecting off a mirror (PF10-03-P01, Thorlabs) (Supplementary Fig. 1a).

Before initiating the thermal stimulation with the IR laser, the visible laser can be positioned over the candidate cells in the culture. The user can then turn the visible alignment laser off and the co-aligned IR laser is modulated with a predefined program of the function generator (DG4162, RIGOL). During these stages, the user can examine the location of the laser and collect the fluorescence emission of cells through the inverted objective below the sample plane, simultaneously. The recorded fluorescence signal is then processed by the proprietary software of the microscope for evaluation.

### Optical imaging and high-precision local laser heating

This imaging setup was used in the experiments where high spatial and temporal resolution is needed, such as heating of single cells or small groups of cells. Optical imaging has been carried out in this study using a home-built upright microscope (Supplementary Fig. 1d,e) integrated, as its unique feature, with a system for high-precision, subcellular-resolution local heating of plant samples with accurately controlled doses of variable-wavelength laser radiation. Optical excitation of genetically encoded fluorescent markers inside sample, collection of their fluorescence response, and laser heating are performed in this microscopy setting via a single 20x, NA = 0.5 microscope objective (MO in Supplementary Fig. 1d) with a working distance of 2.1 mm (Olympus UPlanFLN or equivalent). The fluorescence of genetically encoded markers is driven via two excitation pathways using a two-wavelength source based on two light-emitting diodes D1 (Thorlabs M505L4 or equivalent) and D2 (Thorlabs M565L3 or equivalent). The respective excitation bandwidths are isolated by letting these beams, referred to hereinafter as “blue” and “yellow” light beams, pass through bandpass filters BPF1 (Thorlabs ET500/40x Chroma or equivalent) and BPF2 (Thorlabs MF559-34 or equivalent) to provide excitation of mVenus fluorescent protein, used as a TRP tag and referred to hereinafter as a “green” marker, within the range of 480 to 520 nm and the R-GECO1.1 protein (“red marker), employed as a Ca^2+^ indicator. The blue and yellow beams are combined with a dichroic mirror DM1 (T525lpxr Chroma or equivalent) and DM2 (T525lpxr Chroma, T585lpxr Chroma, or equivalent).

Fluorescent images are recorded with a thermoelectrically cooled, low-noise 4-megapixel CCD camera (Thorlabs4070M-GE-TE or equivalent). Scattered radiation background is suppressed with bandpass and short-pass filters (BPF3 and SPF in Supplementary Fig. 1d). Fluorescence readouts from mVenus and R-GECO1.1 are selected with suitable spectral filters (Thorlabs MF535-22 and Thorlabs T585lpxr Chroma, respectively). The images are recorded with a 4×4 binning, an exposure time of 300 ms, and a frame rate of 3 frames per second.

Local heating is performed in this microscopy setting with trains of near-infrared (NIR) femtosecond laser pulses delivered by a wavelength-tunable optical parametric oscillator (OPO) based on a fan-out periodically poled potassium titanyl phosphate (PPKTP) crystal, synchronously pumped with a 76-MHz, 800-nm, 150-fs, 40-nJ output of a mode-locked Ti: sapphire laser. The signal-wave OPO output has a typical pulse width of 200 fs, pulse energy up to 8 nJ, pulse repetition rate of 76 MHz, and is smoothly tunable within the range of wavelengths from about 970 to 1600 nm. A fast optical beam shutter BSH (Thorlabs SHB05T or equivalent) is used to produce trains of 76-MHz, 200-fs laser pulses with a variable duration, thus providing smoothly adjustable radiation doses for a well-controlled heating of targeted cells. The wavelength tunability of laser radiation proves to be the key for the locality and selectivity of laser heating of targeted cells in bulk samples^20,54^. In experiments presented here, the best results were achieved with the central wavelength of the signal-wave OPO output set within the range of 1300 – 1330 nm. Radiation with longer wavelengths is subject to strong absorption, which makes well-controlled local heating at sufficiently large depths impossible. At shorter wavelengths, on the other hand, absorption is too weak. In this regime, laser powers required for efficient heating often become prohibitively high, leading to an intolerable thermal load on targeted cells^20,55^.

The NIR laser beam used for a local heating of targeted cells enters into the microscope beam path via a dichroic mirror DM3 (Thorlabs DMSP950R or equivalent) and is focused into a sample with the same 20x, NA = 0.5 microscope objective (MO in Supplementary Fig. 1d). Chromatic aberrations of the NIR beam are corrected with an adjustable home-built telescope (TS in Supplementary Fig. 1d). The size of the NIR beam on the targeted site of laser heating can be controlled with an accuracy well within 1.5 μm, with the diameter of this beam typically set at about 3 μm. The background temperature within the samples is stabilized with a suitable temperature controller (Bioscience Tools TC-1-100 or equivalent) and read out with a *negative temperature coefficient* thermistor (NCT in Supplementary Fig. 1d).

### Thermogenetic stimulation using PCR machine

An FDA-approved library was kindly provided by the Chemical Facility of EMBL Heidelberg in 10 96-well plates containing 10 µL of 100 µM solutions of individual drugs in each well.

MIN6 cells were seeded in poly-D-lysine coated wells of 96-well PCR plate and transfected with GECCO-P2A-tdTomato to achieve equimolar expression of GECCO and tdTomato. GECCO expression was normalized for each well by tdTomato fluorescence intensity recorded with a plate reader. Before the experiment, wells were washed with the fresh medium to remove already secreted insulin. After 30 minutes incubation 10 μl of medium were sampled for control measurement (baseline insulin secretion level) and replaced with 10 μl of the fresh medium. Baseline insulin secretion mean value was used to exclude outlier wells from the further consideration.

To find the optimal stimulation pattern, various PCR machine programs were tested. 40 minutes after the program started, the medium of each well was sampled for ELISA insulin measurement.

A for their ability to affect GECCO-induced insulin release. Cells were pre-incubated with a library of FDA-approved drugs drugs for 30 min before the stimulation and subjected to the optimal stimulation protocol. Drug effects on insulin release were calculated with the following formula:

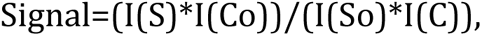

where I(So) is the insulin concentration in the drug treated sample before stimulation, I(S) – the insulin concentration in the drug treated sample after stimulation, I(Co) – the insulin concentration in the vehicle treated sample (MEM medium) before stimulation, I(C) – the insulin concentration in the vehicle treated sample (MEM medium) after stimulation

### Normalization of the channel expression and transfection levels

A Tecan infinite m1000 Pro plate reader equipped with a mCherry filter was used for recording MIN6 transfection levels in 96-well plates. Measurements were performed in microscope imaging solution (115 mM NaCl, 1.2 mM CaCl_2_, 1.2 mM MgCl_2_, 1.2 mM K_2_HPO_4_, 18 mM HEPES, 0-20 mM glucose, pH 7.2). tdTomato labeling was imaged in fluorescence mode (bottom IF, lag time: 0 µs, integration time: 40 µs, flashes: 50, z position calculated from well, settle time: 0, at 37°C and 85% humidity. Excitation wavelength channel: 550(20) nm and emission wavelength channel: 650 (20) nm were used. Wells with significantly lower levels of tdTomato signal were excluded from the comparison. (0-4 wells per 96-well plate).

### Generation, growth and analysis of plant material

Rattle snake (*Crotalus atrox*) caTRPA1 coding sequence (GenBank: GU562967.1) was Arabidopsis codon optimized and synthesized in three Golden (Green) Gate compatible GeneArtTM StringsTM DNA Fragments (Thermo Fisher Scientific). The DNA fragments were subcloned into pJET using the CloneJET PCR Cloning Kit (Thermo Fisher Scientific) and confirmed by sequencing (eurofinsgenomics.eu). The three fragments were then ligated together into pGGC000 via GreenGate cloning^44^ resulting in pGGC-TRPA1CaR. The plant expression vector was assembled via GreenGate cloning using the modules pGGA-pAPA1 (1970 bp promoter fragment of *Arabidopsis thaliana* gene AT1G11910 harboring point mutations to remove two BsaI/Eco31I restriction sites), pGGB003 (N-decoy), pGGC-TRPA1CaR, pGGD-GSL-mTurquoise, pGGE-tHSP18.2M, pGGF-BarR and pGGZ003^44,47,56^ resulting in the pGGZ-RW173 plasmid. *Agrobacterium tumefaciens* strain ASE containing the pSOUP helper plasmid was transformed with pGGZ-RW173 and used for transformation of *Arabidopsis thaliana* R-GECO1 #7^56^ by floral dip^57^.

Seeds of transformed plants were surface sterilized for 10-15 min in 70 % EtOH, washed 3-4 times in 100 % EtOH and sowed on 0.5 MS media (Duchefa) supplemented with 5 mM MES-KOH pH 5.8, 0.8% phytoagar and 10 mg mL^−1^ Glufosinate-ammonium (Fluka) for herbicide selection. After 3-6 days of stratification in the dark at 4°C, transgenic plants were grown for six-days in a growth room (16 h day/8 h night, 22°C, 65% relative humidity, photon fluence rate 100 µmol m^−2^ s^−1^). Positive transformants were transferred to herbicide free 0.5 MS media-containing petri dishes, and on the next day expression of caTRPA1-GSL-mTurquoise (mTurquoise fluorscence emission) was confirmed by visual inspection at a Zeiss Discovery.V20 fluorescence stereo microscope equipped with a GFP filter and a Plan S 1.0x FWD 81 mm lens. At least 40 herbicide resistant and fluorescing seedlings were transferred to round 7 cm pots containing classic soil (Einheitserde) and grown until seed ripening in a growth chamber (Percival I-36LLVL) at (16 h day/8 h night, 17°C, 80% relative humidity, photon fluence rate 50-60 µmol m^−2^ s^−1^). Note that mature caTRPA1 expressing plants do not survive at 22°C. Three independent alleles of R-GECO1/RW173 (pAPA1:caTRPA1-GSL-mTurquoise) have been generated. However, only allele R-GECO1/RW173_20-9 has been used for thermogenetics experiments.

For thermogenetics experiments, T4 seeds of R-GECO1/RW173_20-9 were sown in four horizontal rows on square petri dishes containing 0.5 MS media supplemented with 5 mM MES-KOH pH 5.8 and 0.8% phytoagar, stratified for six days in the dark at 4°C and grown for 4 days in a growth room. Before imaging, seedlings were transferred to microscope dishes (D35-14-0-N, Cellvis) supplemented with 0.25 MS media, 10 mM MES-Tris pH 5.6, 0.7% low melting point agarose (Roth) and topped with imaging buffer (5 mM KCl, 50 µM CaCl_2_, 10 mM MES-Tris pH 5.6).

10 mM MgATP stock (Sigma) dissolved in imaging buffer and pH adjusted with Tris to 5.6 was used for extracellular ATP treatments.

Image processing and analyses were conducted using Fiji^58^. Image processing included background subtraction (∼200), Gaussian blur (1), median (1), 32-bit conversion and thresholding (∼20) to remove background noise. Normalized fluorescence intensity analyses of R-GECO1 (ΔF/F_0_) and vertical profile response maps were generated using a custom build GEFI analyzer Fiji plugin (written by Christian Waadt)^47^. Average response maps were generated using the Z projection average intensity projection type in Fiji. Graphs depicting means ± SD responses were assembled in OriginPro and final figures were assembled in Adobe Illustrator.

## Authors’ contribution

YGE, RW, AMZ, CS and VVB conceived the project; RW generated the transgenic Arabidopsis plants, contributed to experimental design and data analyses of GreenGECCO experiments; MSO, RP, AAL, AC, MP and AMZ designed and built the optical systems for the thermogenetic stimulation and contributed to thermogenetics experiments; VP, IK, DS contributed to genes design and cloning, and fluorescent microscopy; AMM assisted with insulin measurements; CT assisted with microscopy and image analysis; KK performed insulin measurements; PMB and ESN contributed to imaging experiments; YGE, RW, KS, AMZ, RP, CS and VVB analyzed and discussed the data; YGE, RW, MSO, AMZ, CS and VVB and wrote the manuscript

## Notes

### Competing Interest Statement

The authors have declared no competing interest.

