## Supplementary information for "Thermogenetic control of Ca^2+^ levels in cells and tissues"

### Supplementary Notes

#### Cheracterization of GreenGECCO performance in plants

To initially test the utility of GreenGECCO in plant systems, we transiently transformed tobacco *Nicotiana benthamiana* leaves with a mixture of agrobacteria *Agrobacterium tumefaciens,* transfected with plasmids encoding GreenGECCO-P2A-mVenus (Supplementary Fig. 3a) and the fluorescent calcium probe R-GECO1, and performed NIR laser stimulation. Activation of GreenGECCO resulted in a cytosolic calcium burst in epidermal cells within ~1 sec from the moment of radiation supply. From the activated region, the cytosolic $Ca^{2+}$ signal radially propagated to neighboring cells, and after the cessation of the stimulation, the $Ca^{2+}$ signal decreased within 40-50 s (Supplementary Fig. 3b). The stimulation cycle could be repeated several times (Supplementary Fig. 3c). As the width of the laser beam (10 μm) was less than the linear size of the plant cell (30-40 μm), we assume that a single cell $Ca^{2+}$ transient can induce the calcium wave spreading through the tissue

#### $Ca^{2+}$ propagates between plant organs

To better understand the spatial control of thermogenetic stimulation in plants, we excited regions of the root and the hypocotyl in the same field of view, and analyzed the propagation of the induced $Ca^{2+}$ signals. $Ca^{2+}$ waves excited in either the hypocotyl or the root propagated from their origin to neighboring tissues. Signals originating in the hypocotyl were transmitted more efficiently to the root than vice versa. Moreover, the $Ca^{2+}$ waves propagated more efficient in the excited tissues, indicating that the root-hypocotyl junction limits the $Ca^{2+}$ propagation to some extent. Nevertheless, $Ca^{2+}$ waves were efficiently transmitted between roots and areal tissues, indicative that plants can function as a single signaling system (Supplementary Fig. 4d,e).

### Supplementary Figures

#### Supplementary Fig. 1: The NIR thermal stimulation modules.

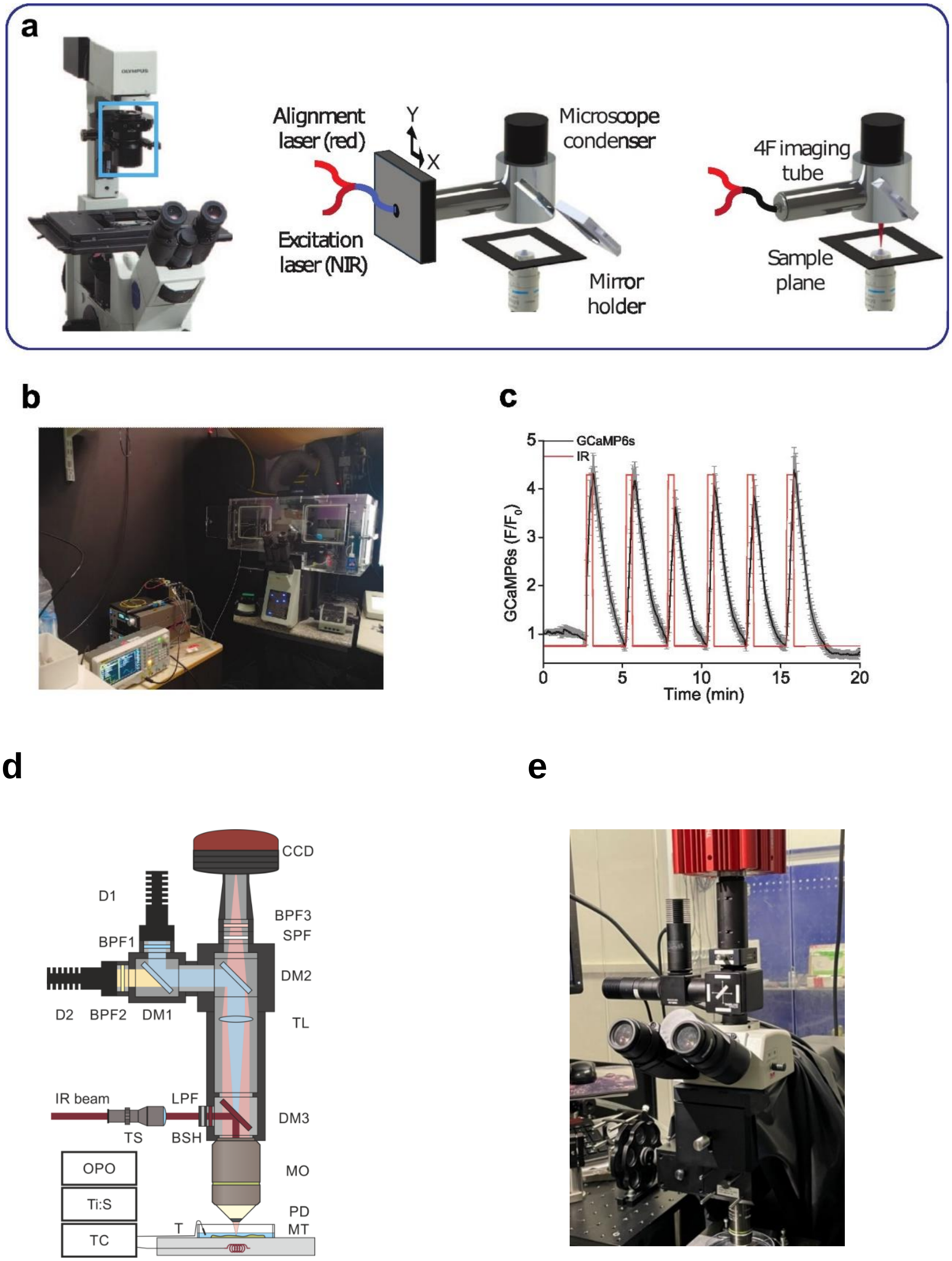

**a** Left image: The module for heating of large groups of cells can fit any inverted microscope setup by fitting the module head into the condenser (blue rectangle). Central image: NIR and visible red laser combined to guide the NIR focal spot on the sample. The distal end of the fiber was connected to an x-y stage to align the laser to the optical axis. Right image: Mirror holder can be slided in/out to swap between epi-configuration bright field imaging and NIR stimulation mode, seamlessly. **b** A photograph of the module **a**. **c**, Fluorescence intensity of GCaMP6s (black line) reflects the $Ca^{2+}$ response in HEK293 cells to repetitive 148 mW laser stimulations (red line). **d,e** A schematic **d** and a photograph **e** of the microscopy setting, enabling optical imaging simultaneously with a subcellular-resolution, high-precision heating: Ti:S, mode-locked Ti: sapphire; OPO, optical parametric oscillator; D1, D2, light-emitting diodes; IR beam, near-infrared laser beam; BPF1, BPF2, BPF3, bandpass filters; DM1, DM2, DM3, dichroic mirrors; SPF, shortpass filter; LPF, longpass filter; TL, tube lens; TS, telescope; BSH, beam shutter; MO, microscope objective; PD, Petri dish; MT, microscope table; T, NCT, thermistor; TC, temperature controller; CCD, CCD camera.

**Supplementary Fig. 2: Thermogenetic control of $Ca^{2+}$ oscillations in functionally coupled MIN6 cells.**

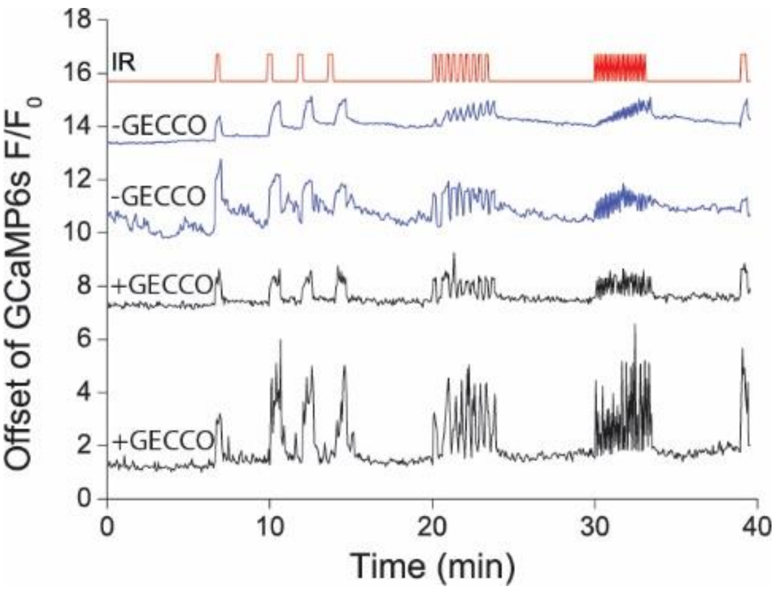


Combination of different train frequencies and durations (3-100 sec, 0.5-10 Hz) at a constant power of 58 mW induce similar Ca2+ responses in cells expressing GECCO and non-transfected cells within the same cell culture.

**Supplementary Fig. 3: Themogenetic $Ca^{2+}$ control in *Nicotiana benthamiana.***

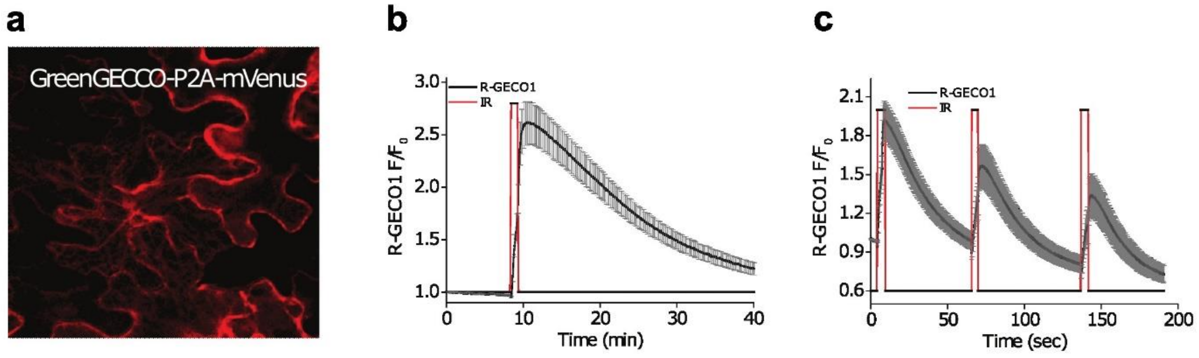


**a** *N. benthamiana* leaves transfected with plasmid encoding GreenGECCO-P2A-mVenus. **b,c** $Ca^{2+}$ imaging in leaves controlled by10 μm wide laser beam at 100 mW nominal radiation power.

**Supplementary Fig. 4: Control experiments, stimulations in the root-hypocotyl-junction and subcellular localization of GreenGECCO.**

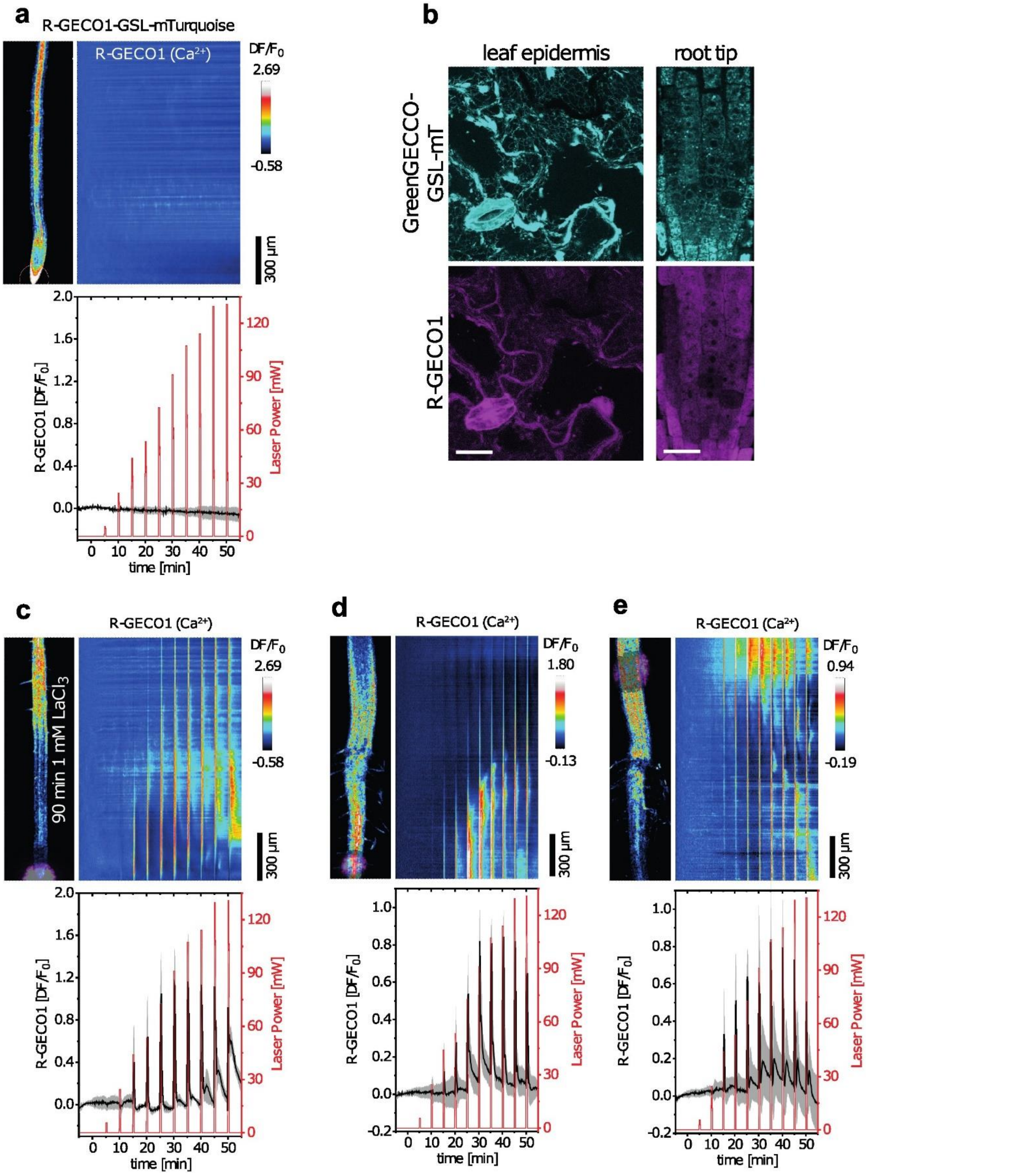

**a, c-e** Four-day-old Arabidopsis seedlings, expressing **a** R-GECO1-GSL-mTurquoise or **c-e** caTRPA1-GSL-(mT)urquoise (GreenGECCO) in the R-GECO1 $Ca^{2+}$ probe background, were thermogenetically stimulated with 30 sec pulses of increasing NIR laser power with 4.5 min gaps between each pulse. NIR laser stimulations were applied to **a and c** root tips (means ± SD, n = 4-5), **d** upper root regions (means ± SD, n = 6) or **e** hypocotyls (means ± SD, n = 5). Experiments in **c** were conducted after 90 min pretreatment with 1 mM of the $Ca^{2+}$ channel blocker $LaCl_3$. The left images indicate the respective R-GECO1 fluorescence intensity at t = -5 min, with the purple spot **c-e** or white circle in **a** representing the regions of NIR laser stimulation. The right images indicate normalized average spatiotemporal response profiles of R-GECO1 (color scale bar on the right) along the root axis with the same time scale as in the graph below (frame rate 10 $min^{-1}$). The graphs indicate normalized average R-GECO1 responses (black lines), calculated from whole image mean values, and the applied NIR laser power (red lines). **b** Confocal laser scanning microscope (CLSM) analyses of the subcellular localization of caTRPA1-GSL-(mT)urquoise (top) and R-GECO1 (bottom) in leaf epidermis cells (left) shown as z-stack maximum projections, and root tip cells (right). The net-like structures of the mTurquoise fluorescence in the leaf image indicate ER membrane localization, whereas diffuse fluorescence patterns of R-GECO1 indicate cytosolic and nuclear localization. Scale bar is 20 μm.

**Table 1: Drugs that induced insulin release after addition at 10 μM concentration but also enhanced thermogenetic insulin release**

| Number | Drug name | Mean response signal factor (ELISA2/1) | SEM of the response |
|---|---|---|---|
| 1 | Rosiglitazone maleate | 2.225 | 0.0854 |
| 2 | Ticlopidine hydrochloride | 2.122 | 0.0808 |
| 3 | Delta1-hydrocortisone 21-hemisuccinate sodium salt | 1.757 | 0.0785 |
| 4 | Altanserin | 2.143 | 0.0776 |
| 5 | Diazoxide | 1.501 | 0.073 |
| 6 | Nialamide | 1.458 | 0.0933 |

| | | | |
|---|---|---|---|
| 7 | Methanesulfonamide | 1.613 | 0.0707 |
| 8 | Omeprazole | 1.774 | 0.092 |
| 9 | Homoharringtonine | 1.938 | 0.0785 |
| 10 | Clobenpropit | 2.218 | 0.0861 |
| 11 | Parecoxib sodium | 1.566 | 0.0813 |
| 12 | Miconazole nitrate | 1.516 | 0.0871 |
| 13 | Piperacillin sodium | 1.54 | 0.0965 |
| 14 | L-thyroxine | 1.934 | 0.0868 |
| 15 | Atropine | 1.826 | 0.0941 |
| 16 | Chlorambucil | 1.458 | 0.093 |
| 17 | Oxybutynin chloride | 1.889 | 0.0809 |
| 18 | Medroxyprogesterone 17-acetate | 1.666 | 0.0784 |
| 19 | Pyrimethamine | 1.459 | 0.0794 |
| 20 | Hydrocortisone hemisuccinate | 1.317 | 0.0752 |
| 21 | Pro-banthine | 2.013 | 0.0934 |
| 22 | Spectinomycin dihydrochloride pentahydrate | 1.339 | 0.0869 |

| | | | |
|---|---|---|---|
| 23 | Westcort | 1.45 | 0.0796 |
| 24 | Selegiline hydrochloride | 1.804 | 0.0741 |
| 25 | Nafcillin sodium salt monohydrate | 2.04 | 0.0822 |
| 26 | Tolazamide | 1.488 | 0.0737 |
| 27 | Dibenzyline (=Phenoxybenzamine hydrochloride) | 1.356 | 0.0792 |
| 28 | Ethambutol | 1.557 | 0.0865 |
| 29 | Clopidogrel | 1.402 | 0.0908 |
| 30 | Quinapril hydrochloride | 1.521 | 0.0876 |
| 31 | Citalopram | 2.039 | 0.0739 |
| 32 | Simvastatin | 2.17 | 0.0858 |
| 33 | HMBA (N,N′-Hexamethylene bis(acetamide)) | 1.856 | 0.0723 |
| 34 | Bifemelane | 1.521 | 0.0794 |
| 35 | Mesalamine | 2.178 | 0.092 |
| 36 | Galanthamine | 1.875 | 0.0863 |
| 37 | Aurorix | 1.535 | 0.0966 |

| | | | |
|---|---|---|---|
| 38 | Kitasamycin | 2.168 | 0.0881 |

**Table 2. Drugs that by themselves did not induce insulin release after addition at 10 µM concentration but enhanced thermogenetic insulin release.**

| Number | Drug name | Mean response signal factor (ELISA2/1) | SEM of the response |
|---|---|---|---|
| 1 | Escitalopram oxalate | 1.748 | 0.0755 |
| 2 | Oligomycin c | 1.557 | 0.073 |
| 3 | Moxonidine hcl | 1.61 | 0.0817 |
| 4 | Modafinil | 1.971 | 0.0814 |
| 5 | Procarbazine hydrochloride | 1.303 | 0.0811 |
| 6 | 2-chloroadenosine | 2.123 | 0.096 |
| 7 | Exemestane | 1.988 | 0.0712 |
| 8 | Dexchlorpheniramine maleate | 2.173 | 0.0757 |
| 9 | Tramadol | 1.916 | 0.0933 |
| 10 | Homoveratrylamine | 2.073 | 0.0894 |
| 11 | Dehydrocholic acid | 1.724 | 0.0977 |

| | | | |
|---|---|---|---|
| 12 | Vindesine sulfate | 1.339 | 0.0992 |
| 13 | Glimepiride | 1.492 | 0.0835 |
| 14 | Pyridine-2-aldoxime methochloride | 1.751 | 0.0938 |
| 15 | Sulfinpyrazone | 1.889 | 0.08 |
| 16 | Labetalol hydrochloride | 1.83 | 0.0785 |
| 17 | Nabumetone | 1.831 | 0.0992 |
| 18 | Phylloquinone | 1.888 | 0.0903 |
| 19 | Memantine hydrochloride | 1.309 | 0.0702 |
| 20 | Dicloxacillin sodium | 2.015 | 0.0864 |
| 21 | Rifabutin | 1.568 | 0.0983 |
| 22 | Alprazolam | 1.963 | 0.0926 |
| 23 | Secnidazole | 2.177 | 0.0856 |
| 24 | Stavudine | 1.501 | 0.0968 |
